# Sleep and circadian rhythm activity alterations during adolescence in a mouse model of neonatal fentanyl withdrawal syndrome

**DOI:** 10.1101/2024.07.05.602239

**Authors:** Benjamin R. Williams, Mackenzie C. Gamble, Navsharan Singh, Camron D. Bryant, Beth A. Logan, Ryan W. Logan

## Abstract

**Purpose:** Fentanyl, a highly potent synthetic opioid, is a major contributor to the ongoing opioid epidemic. During adulthood, fentanyl is known to induce pronounced sleep and circadian disturbances during use and withdrawal. Children exposed to opioids *in utero* are likely to develop neonatal opioid withdrawal syndrome, and display sleep disturbances after birth. However, it is currently unknown how neonatal opioid withdrawal from fentanyl impacts sleep and circadian rhythms in mice later in life.

**Methods:** To model neonatal opioid withdrawal syndrome, mice were treated with fentanyl from postnatal days 1 through 14, analogous to the third trimester of human gestation. After weaning, fentanyl and saline treated mice underwent non-invasive sleep and circadian rhythm monitoring during adolescence postnatal days 23 through 30.

**Results:** Neonatal fentanyl exposure led to reduced duration of wake and a decrease in the number of bouts of non-rapid eye movement sleep. Further, neonatally exposed mice displayed an increase in the average duration of rapid eye movement sleep bouts, reflecting an overall increase in the percent time spent in rapid eye movement sleep across days.

**Conclusions:** Neonatal fentanyl exposure leads to altered sleep-wake states during adolescence in mice.

## INTRODUCTION

An estimated 8 million Americans are currently living with opioid use disorder (OUD), with opioid overdoses accounting for nearly 70,000 deaths annually (Keyes et al, 2022). In recent years, fentanyl has become particularly prevalent in cases of opioid misuse and overdose. In addition to the risk of overdose, chronic opioid use is associated with numerous side effects including severe sleep and circadian disruptions (Korff et al, 2012 ; Xue et al, 2022). Opioid withdrawal-related sleep disturbances are of particular concern as they are known risk factors for craving and relapse, further increasing the difficulty of achieving abstinence (Brower & Perron, 2010 ; Roehrs & Roth, 2015).

Notably, in pregnant mothers, the negative effects of long-term opioid use are heightened, as opioids are known to cross the placental barrier, posing the risk of *in utero* opioid exposure. In as many as 94% of prenatal opioid exposure cases, neonates are born experiencing opioid withdrawal in a condition known as neonatal opioid withdrawal syndrome (NOWS) (Zanckl et al., 2021), particularly when exposure occurs during the third trimester of gestation (Desai et al., 2015). Neonates suffering from opioid withdrawal may present with low weight, hyperirritability, microcephaly, or more serious complications including respiratory difficulties, seizures, and tremors (Auger et al., 2018 ; Lust et al., 2022 ; Mascarenhas et al., 2024). Much like how opioid-induced sleep and circadian disturbances are extensively reported in adult patients experiencing withdrawal, clinical reports indicate that prenatal opioid exposure disrupts sleep quality in newborns. Specifically, preliminary evidence suggests that opioid exposed newborns experience decreased and more fragmented sleep, compared to unexposed, otherwise, typical newborns (OBrien & Jeffery, 2002). The evidence-based care tool known as “Eat, Sleep, Console”, considers an infant’s sleep quality (*i.e*., ability to sleep for one hour undisturbed), as a core metric of opioid withdrawal symptoms and is often used to monitor progression of effective treatment.

Further research is needed to characterize how neonatal opioid withdrawal syndrome impacts sleep-wake states (*i.e*. wake, non-rapid eye movement sleep, rapid eye movement sleep). In the present study, we used Carworth Farms White (CFW) mice, an outbred mouse population, to model neonatal opioid withdrawal syndrome from fentanyl (Borrelli et al., 2021 ; Borrelli et al., 2023) and investigate the impact of fentanyl exposure during early development on sleep and circadian rhythms during adolescence.

## MATERIALS AND METHODS

### Animals

Breeder pairs of Carworth Farms White (CFW) mice (Charles River Laboratories; Shrewsbury, MA, USA; IMSR Cat# Crl:CFW(SW); Facility Strain Code: 024), ages 8-12 weeks, were maintained on a 12:12 reverse light-dark cycle (lights on at 22:00, zeitgeber time (ZT) 0, and lights off at 10:00, ZT12). At birth, male and female mice were randomly assigned to drug treatment groups. Food and water were provided *ad libitum*. Animal use and procedures were conducted in accordance with the National Institute of Health guidelines and approved by the Boston University School of Medicine Institutional Animal Care and Use Committee.

### Drug Administration

Fentanyl citrate powder was purchased from Covetrus and dissolved with 0.9% saline (Fisher Scientific). Mice were randomly assigned treatment groups at birth and injected subcutaneously (s.c.) twice daily for 14 days (P1 to P14) with either saline (10 ml/kg), or fentanyl (0.1 mg/kg at 10 ml/kg). Drugs were administered between ZT9-10, and ZT18-19.

### Sleep & Circadian Monitoring

From postnatal day (P) 23 to P30, mice underwent non-invasive three-state sleep-wake analysis using PiezoSleep Mouse Behavioral Tracking Systems (Signal Solutions, Inc., Lexington, KY, USA). For the duration of the sleep recording, mice were single housed with food and water provided *ad libitum*, under identical reverse light-dark cycle conditions within sound attenuating and light controlled cabinets (Tecniplast; Westchester, PA, USA). Mice were allowed to habituate to the Piezo sleep boxes for 2 days prior to data acquisition.

### Experimental Design

### Statistical Analysis

Hourly or daily data was analyzed as two-way repeated measures ANOVA (GraphPad Prism version 10.1). Averaged summary data was assessed for normality using a Shapiro-Wilk test, then analyzed as two sample t-tests (Mann-Whitney U for nonparametric measures) with Welch’s correction applied when F test to compare variances failed (GraphPad Prism version 10.1). Data is presented as mean ± standard error of mean with significance set at □ = 0.05. All analyses are presented as combined data from male and female mice as no sex differences were detected.

## RESULTS

### Impact of neonatal fentanyl exposure in mice on diurnal activity in adolescence

After birth, at P1, mice were administered fentanyl (0.1mg/kg) subcutaneously twice daily for 14 days to model neonatal opioid withdrawal syndrome (Borrelli et al., 2021 ; Borrelli et al., 2023). Locomotor activity, respiration and heart rates, as part of the PiezoSleep system, were recorded continuously for 8 days beginning after weaning at P21. To investigate the impact of neonatal fentanyl exposure on diurnal rhythms in adolescence, we compared diurnal locomotor activity on P23-30 between saline and fentanyl exposed mice. Activity bouts were assessed in 6-minute bins and averaged across saline or fentanyl groups for days P23-30. As expected, the average diurnal activity was greater during the dark phase of light-dark cycle in saline mice (Figure 2A). Fentanyl exposed mice displayed a similar diurnal activity pattern (Figure 2A).

**Figure 1.**
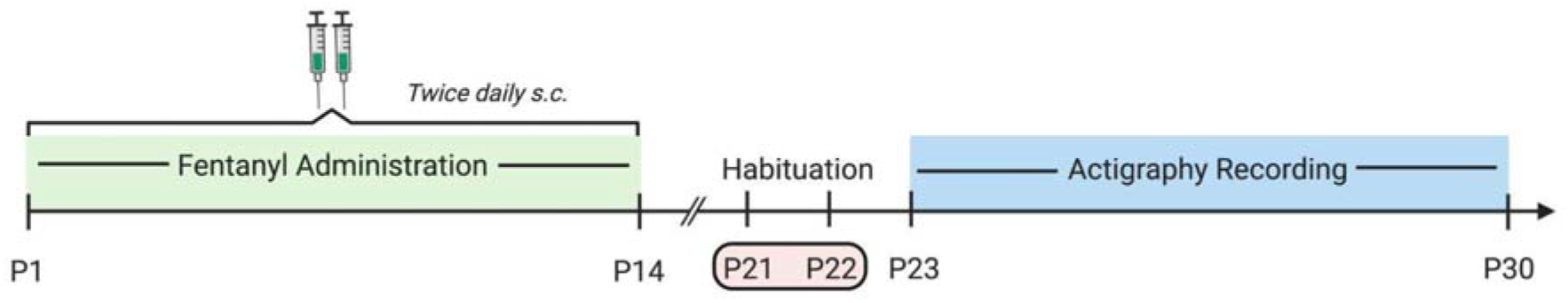
Schematic of experimental timeline. Male and female Carworth Farms White mice were administered fentanyl (s.c.) twice daily from P1 to P14. After weaning, at P21, mice weretransferred to PiezoSleep chambers and habituated for 2 days before undergoing continuous actigraphy recording for 8 days (P23-P30).

**Figure 2.**
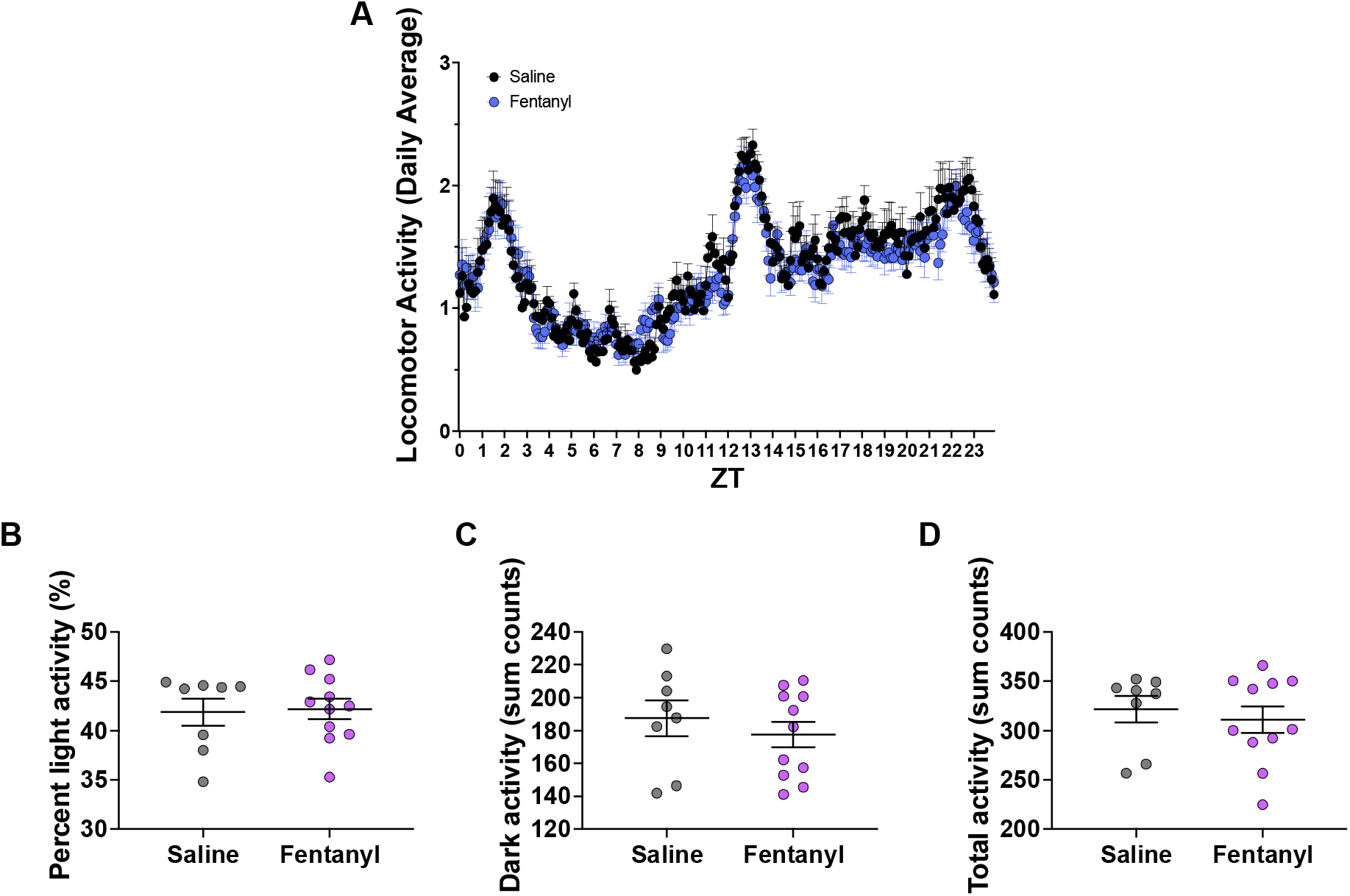
Diurnal activity during days P23-30. (A) Hourly diurnal activity of saline (n=4 male, 4 female) and fentanyl (n=6 male, 5 female) mice averaged in 6 min bins, across days P23-30 (B) Percent of activity occurring during the day (light period) averaged across days P23-30. (C) Sum of activity counts occurring during the night (dark period), averaged across days P23-30.(D) Sum of activity counts occurring during a 24-hr period averaged across days P23-30. Error bars represent mean ± SEM.

There was no significant difference observed in the average sum of activity counts occurring in the light period across all days between saline (M = 41.867 ± 1.372) and fentanyl (M = 42.194 ± 1.037; P = 0.968) treated mice (Figure 2B), or in the average number of night activity counts (Saline: M = 187.502 ± 10.816, Fentanyl: M = 177.565 ± 7.933; P = 0.458) (Figure 2C).Similarly, no significant difference in total activity accounts between saline (M = 321.838 ± 13.455) and fentanyl (M = 311.031 ± 13.450; P = 0.840) treated mice was observed when averaged across a 24-hour period (Figure 2D). Thus, neonatal fentanyl administration did not impact diurnal activity rhythms in mice during adolescence.

### Neonatal fentanyl exposure led to decreased non-rapid eye movement sleep and increased rapid eye movement sleep bout durations

We also investigated the impact of neonatal fentanyl exposure on sleep-wake states in mice during adolescence. Both saline and fentanyl exposure mice spent most of their time in wake, followed by NREMS and REMS. No significant difference in average wake bout duration on days P23-30 was observed between saline and fentanyl treated mice (Drug: F_(1, 17)_ = 0.199, P = 0.662, Time: F_(7, 119)_ = 0.975, P = 0.453, Interaction: F_(7, 119)_ = 0.905, P = 0.505) (Figure 3A), though fentanyl treated mice displayed a decreased average wake bout duration when data across all recording days was pooled (Saline: M = 33.569 ± 0.290, Fentanyl: M = 31.927 ± 0.484, P = 0.011) (Figure 4A). When assessing the average percent of time spent awake across days P23-30, a significant main effect was observed for time (F_(7, 119)_ = 6.100, P < 0.001) but not for drug (F_(1, 17)_ = 0.035, P = 0.854), interaction (F_(7, 119)_ = 1.112, P = 0.360) (Figure 3B), or when averaged across all days of recording (Saline: M = 49.087 ± 0.351, Fentanyl: M = 48.699 ± 0.557, P = 0.798) (Figure 4B). Similar results were observed when analyzing average NREMS bout duration for days P23-30 in which a significant main effect was found for time (F_7, 119)_ = 8.508, P < 0.001) but not for drug (F_(1, 17)_ = 1.647, P = 0.217), or interaction (F_(7, 119)_ = 1.350, P = 0.233) (Figure 3C). When averaged across all days of recording, fentanyl treated mice displayed a significantly shorter average NREMS bout duration compared to saline controls (Saline: M = 18.078 ± 0.258, Fentanyl: M = 16.651 ± 0.284, P = 0.002) (Figure 4C). While neonatal fentanyl treatment decreased the average NREMS bout duration across all days, it did not impact the average percent of time spent in NREMS on individual days P23-30 (Time: F_(7, 119)_ = 3.277, P = 0.003, Drug: F_(1, 17)_ = 0.465, P = 0.504, Interaction: F_7, 119)_ = 1.711, P = 0.113 (Figure 3D). However, pooled data revealed that fentanyl treated mice displayed decreased percent NREMS (Saline: M = 42.304 ± 0.242, Fentanyl: M = 40.922 ± 0.375, P = 0.008) (Figure 4D). Similar to wake and NREMS, fentanyl treatment did not impact the average REMS bout duration during days P23-30 (Drug: F_1, 17)_ = 0.880, P = 0.361), Time: F_(7, 119)_ = 1.127, P = 0.351, Interaction: F_(7, 119)_ = 1.173, P = 0.324) (Figure 3E), nor the average percent of time spent in REMS during days P23-30 (Time: F_(7, 119)_ = 15.39, P < 0.001, Drug: F_(1, 17)_ = 3.318, P = 0.086, Interaction: F_(7, 119)_ = 0.973, P = 0.455) (Figure 3F). Notably, when averaged across all days of recording, fentanyl treatment significantly increased the average REMS bout duration (Saline: M = 6.390 ± 0.051, Fentanyl: M = 6.596 ± 0.056, P = 0.016) (Figure 4E) as well as the average percent of time spent in REMS (Saline: M = 8.609 ± 0.418, Fentanyl: 10.517 ± 0.473, P = 0.009) (Figure 4F). Taken together, these findings suggest that, over extended actigraphy recordings, neonatal opioid exposure increases average wake bout duration, decreases both the average bout duration and percent time in NREMS, and increases both the average REMS bout duration and percent of time spent in REMS.

**Figure 3.**
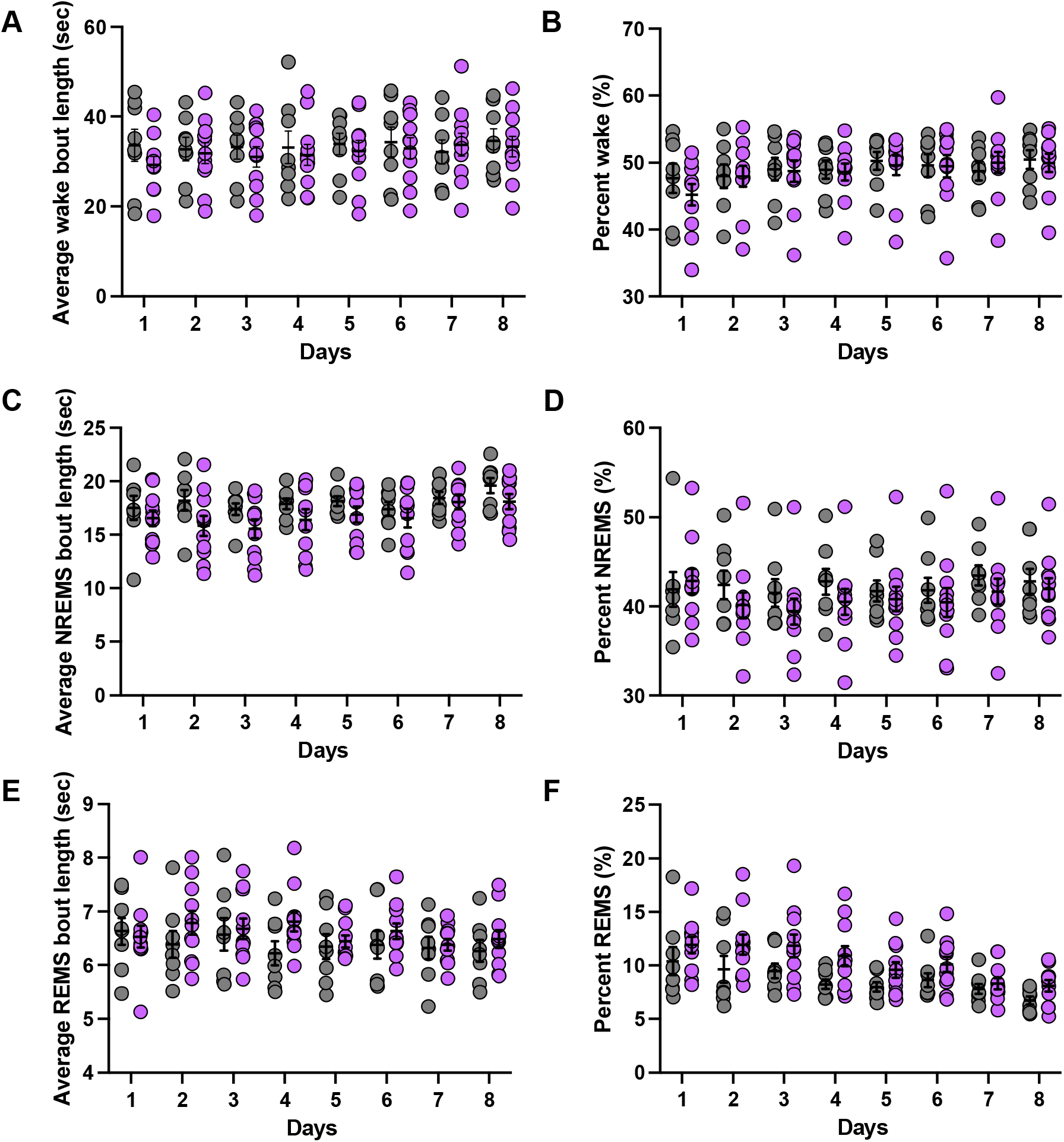
Sleep-wake composition during days P23-30. (A) Average duration of wake bouts (sec) occurring daily. (B) Average percent of time spent in wake occurring daily. (C) Average duration of NREMS bouts (sec) occurring daily. (D) Average percent of time spent in NREMS occurring daily. (E) Average duration of REMS bouts (sec) occurring daily. (F) Average percent of time spent in REMS occurring daily. Error bars represent mean ± SEM.

**Figure 4.**
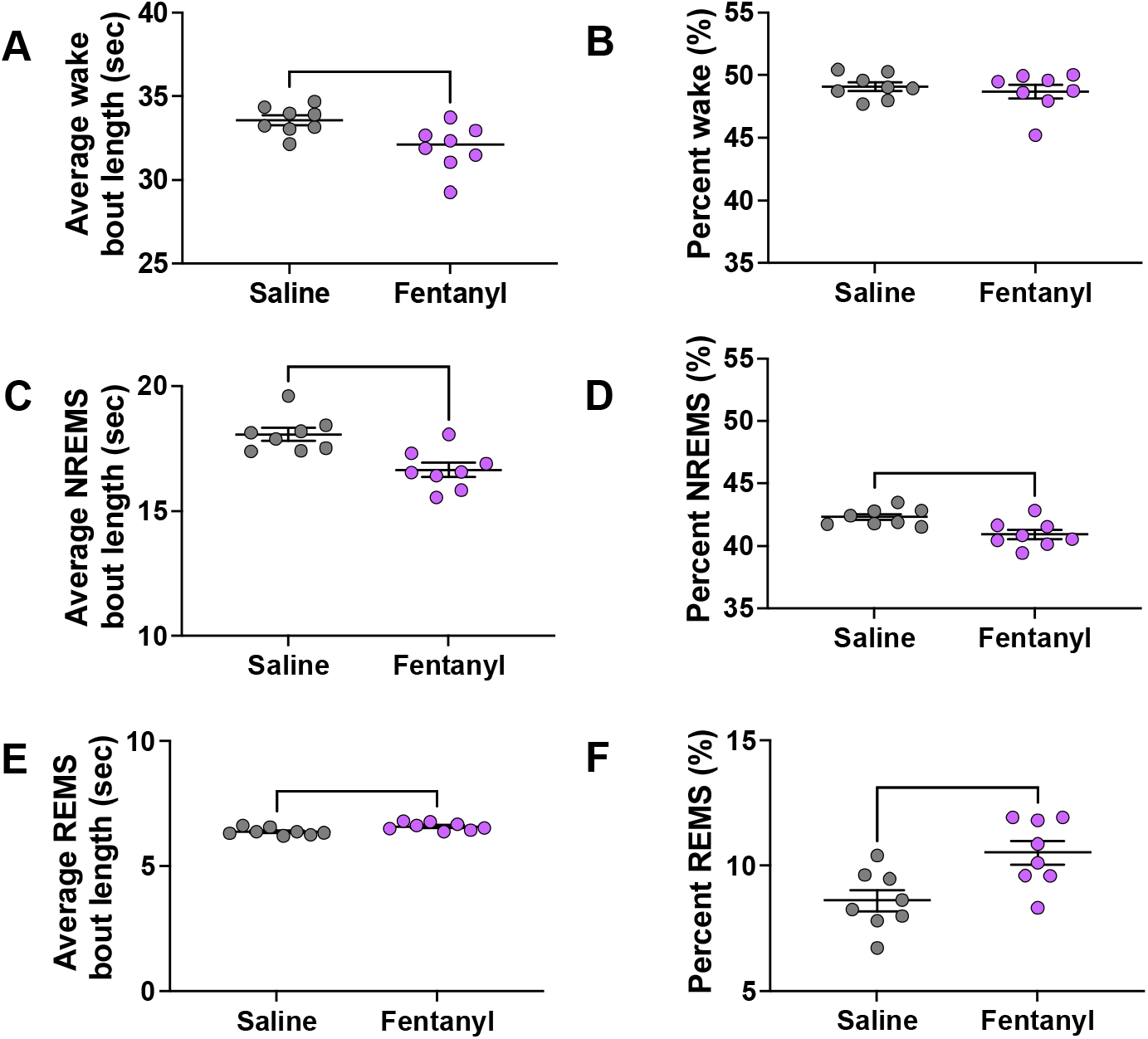
Sleep-wake composition averaged across all days. (A) Average duration of wake bouts (sec). (B) Average percent of time spent in wake. (C) Average duration of NREMS bouts (sec). (D) Average percent of time spent in NREMS. (E) Average duration of REMS bouts (sec).(F) Average percent of time spent in REMS. Data presented as mean ± SEM. * = P ≤ 0.05, ** = P ≤ 0.01, *** = P ≤ 0.001.

## DISCUSSION

Chronic opioid use and misuse persists as a public health crisis, with fentanyl posing a major threat to prolonged use-associated side effects, and potential overdose. Beyond risk to the individual, opioid use during pregnancy has been implicated in long-term developmental impairments collectively defined in a diagnosis of NOWS. While opioid use in adult humans and mice has been shown to induce lasting sleep and circadian disturbances, particularly during withdrawal, little work has been done to characterize the impact of neonatal opioid withdrawal syndrome on sleep and circadian rhythmicity. And, to date, no work has been done to investigate the specific impact of fentanyl on sleep disturbances following postnatal exposure, particularly during adolescence which presents as a vulnerable period for both psychiatric symptom and substance use onset.

In the present study, we observed changes during adolescence in NREMS and REMS in a mouse model of neonatal opioid withdrawal syndrome, independent of changes in circadian activity. We found that twice daily fentanyl injection from P1 to P14 did not disrupt diurnal activity in mice. These findings are consistent with previous clinical studies in humans (Sarfi et al., 2009), as well as in mice exposed to morphine both *in utero* and postnatally (Dunn et al., 2023). Despite the lack of significant circadian behavioral disturbances, the potential impact of neonatal opioid withdrawal on endogenous molecular rhythms should not be overlooked. In one study, adult rats exposed to morphine both *in utero* and as neonates displayed differential rhythmic gene expression in the suprachiasmatic nucleus, the circadian pacemaker of the body, and the liver as well as an altered rhythm of arylalkylamine N-acetyltransferase within the pineal gland, the main driver of melatonin rhythms (Pačesová et al., 2023). Thus, there appears to be a complex relationship between opioid exposure and homeostatic modulators of sleep and arousal states which is sufficient to induce disturbances at the molecular level but not at the behavioral level.

On per day basis, we also found that neonatal fentanyl administration does not impact the average sleep bout duration, or the percent of time spent in wake, NREMS, or REMS. These findings complement a previous study which reported that morphine-treated male mice did not display any difference in average sleep bout durations, and that morphine-treated female mice displayed similar sleep-wake activity to that of saline-treated controls. Of note, this same study found that morphine-treated male mice displayed differential sleep-wake activity across a 36-hour period, and that morphine-treated female mice exhibited longer sleep bouts thereby suggesting sexually dimorphic sleep alterations following gestational and postnatal opioid exposure (Dunn et al., 2023). Future work should aim to further characterize these sex differences in 3-state sleep architecture (*i.e*., wake, NREMS, and REMS). Intriguingly, when analyzed across the entirety of the recording days, we found that neonatal fentanyl administration decreased average NREMS bout durations and the percent of time spent in NREMS and increased average REMS bout durations and the percent of time spent in REMS. The impact of postnatal fentanyl treatment on sleep alterations appears to be subtle and presents only over the course of extended recordings. In addition, the impact of neonatal fentanyl exposure on sleep in adolescent mice appears to be similar to that of fentanyl-induced sleep disturbances in adult mice which also display changes in NREMS.

In previous work, we show that sleep changes in adult mice differ between acute and chronic fentanyl administration, as well as during acute and chronic withdrawal. Adult mice who received a single injection of fentanyl displayed decreased time spent in NREMS and a decreased number of total sleep bouts which persisted into acute withdrawal (24-48 hours post injection) (Gamble et al., 2022). However, adult mice who received chronic fentanyl treatment twice daily for 7 days did not display a decrease in percent NREMS until cessation of drug administration during chronic withdrawal. Of note, these findings in adult mice indicate that fentanyl-induced sleep alterations are distinct between recordings during drug exposure and eventual withdrawal. In the present study, sleep and circadian activity was only examined in the context of prolonged withdrawal due to technical limitations, as no standard technique currently exists to quantify three-state sleep architecture prior to adolescence in mice. Indeed, it would be interesting to determine if sleep disturbances are present in neonates P1-P14 actively being treated with fentanyl, and if any sleep changes found are distinct from those observed following cessation of drug exposure. Further, the long-term impacts of neonatal fentanyl exposure on sleep and circadian rhythmicity throughout development remains unknown. Future work should aim to investigate if NOWS-associated sleep disturbances persist into adulthood, and if these changes in sleep architecture are associated with disease-related phenotypes such as increased drug craving or propensity to reinstatement.

## AUTHOR CONTRIBUTIONS

**C.D.B, N.S, B.A.L**., **and R.W.L**.: Conceptualization; **B.R.W**., **M.C.G**., **and N.S**.: Data curation**; R.W.L**., **B.R.W**., **and M.C.G**.: Formal analysis; **R.W.L**.: Funding acquisition; **B.R.W**., **M.C.G**., **and N.S**.: Investigation; **B.R.W**., **M.C.G**., **B.A.L**., **and R.W.L**.: Manuscript; **R.W.L**., **N.S**.: Methodology; **R.W.L**., **and N.S**.: Project administration; **C.D.B. and R.W.L**.: Supervision; **B.R.W**., **and M.C.G**.: Visualization.

## NOTES

The authors have no conflict of interest, nor competing financial interest to declare.

## ABBREVIATIONS

OUD: opioid use disorder
NOWS: neonatal opioid withdrawal syndrome
REMS: rapid eye movement sleep
NREMS: non-rapid eye movement sleep
ZT: zeitgeber time

